# Joint Instability Causes Catabolic Enzyme Production in Chondrocytes prior to Synovial Cells in Novel Non-Invasive ACL ruptured Mouse Model

**DOI:** 10.1101/2022.05.28.493828

**Authors:** Kei Takahata, Kohei Arakawa, Saaya Enomoto, Yuna Usami, Koyo Nogi, Riku Saitou, Kaichi Ozone, Haruna Takahashi, Moe Yoneno, Takanori Kokubun

## Abstract

**Objective:** The ACL-deficient model helps to clarify the mechanism of knee OA; however, the conventional ACL injury model could have included concurrent onset factors such as direct compression stress to cartilage and subchondral bone. In this study, we established a novel Non-invasive ACL-Ruptured mouse model without concurrent injuries and elucidated the relationship between OA progression and joint instability.

**Design:** We induced the ACL-Rupture non-invasively in twelve-week-old C57BL/6 male mice and evaluated histological, macroscopical, and morphological analysis at 0 days. Next, we created the ACL-R, controlled abnormal tibial translation (CATT), and Sham groups. Then, the joint stability and OA pathophysiology were analyzed at 2, 4, and 8 weeks.

**Results:** No intra-articular injuries, except for ACL rupture, were observed in the ACL-R model. ACL-R mice increased anterior tibial displacement compared to the Sham group (p<0.001, 95% CI [-1.509 to -0.966]) and CATT group (p<0.001, 95% CI [-0.841 to -0.298]) at 8 weeks. All mice in the ACL-R group caused cartilage degeneration. The degree of cartilage degeneration in the ACL-R group was higher than in the CATT group (p=0.006) at 8 weeks. The MMP-3-positive cell rate of chondrocytes increased in the ACL-R group than CATT group from 4 weeks (p=0.043; 95% CI [-28.32 to -0.364]) while that of synovial cells increased at 8 weeks (p=0.031; 95% CI [-23.398 to -1.021]).

**Conclusion:** We successfully established a Non-invasive ACL-R model without intra-articular damage. Our model revealed that chondrocytes might react to abnormal mechanical stress prior to synovial cells while the knee OA onset.

## Introduction

Knee Osteoarthritis (OA) is a representative musculoskeletal disease, and mechanical stress has been attracting attention as a significant pathogenic factor. Under physiological mechanical stress, chondrocytes promote the synthesis of extracellular matrix and maintain homeostasis ^1^. Conversely, abnormal mechanical stress stimulates more upregulation of catabolic enzymes, such as a disintegrin and metalloproteinase with thrombospondin motifs (ADAMTS-4,5) and matrix metalloproteinase (MMP-1,3,13) that cause articular cartilage degeneration^2-4^.

Since the knee joint is a synovial joint sealed by the joint capsule, the intra-articular tissues, such as the articular cartilage and synovial membrane, biologically interact with the synovial fluid ^5-9^. Synovitis is an intra-articular pathology involved in OA that contributes to cartilage degradation through the release of proinflammatory and catabolic products into the synovial fluid ^10^. Some researchers have reported that synovitis occurs in a relatively early OA stage and causes cartilage degeneration ^11, 12^. However, the relationship between these two pathologies under mechanical stress hasn’t been elucidated in detail.

Many OA animal models induced by mechanical stress have been developed, and a representative one is the ACL-transection (ACL-T) model. We recently reported a joint-controlled model, the controlled abnormal tibial translation (CATT) model ^13^, which suppresses anterior-posterior joint instability *in vivo* due to ACL-T. Moreover, we established another joint-controlled model called the controlled abnormal tibial rotation model, which suppresses abnormal joint rotation due to the destabilization of the medial meniscus (DMM) ^14^. Using these models, Arakawa *et al*. revealed that different mechanical stress types have different effects on joint degeneration. In particular, it was suggested that anterior-posterior joint instability in the ACL-T model causes articular cartilage degeneration by inducing chondrocyte hypertrophy, unlike in the CATT model _15_.

In a previous study, we used an ACL-T model with surgical intervention; however, the synovial invasion may disrupt the natural intra-articular environment, leading to unintended adaptive and healing processes. To avoid these problems, many researchers have attempted to establish non-invasive ACL-R models ^16-18^. Christiansen *et al*. reported a mouse model in which the hind limbs were fixed using a device and the tibia was dislocated anteriorly direction to cause ACL rupture. However, conventional non-invasive ACL-R models have focused on the effect of compression stress on OA progression because these models were created by inducing compression force on the knee joint surface. In addition to the induced joint instability due to ACL-R, primary injury of the cartilage, meniscus, and subchondral bone occur possibility.

Therefore, we aimed to establish a novel, non-invasive ACL-R knee OA model that avoids intra-articular tissue injuries, except for ACL ruptures. We manually applied anterior tibial translation stress to rupture the ACL without any contact between the tibial plateau and the femoral condyle. By comparing this model with the CATT model, we attempted to reveal the onset mechanism of knee OA induced by joint instability, especially in synovitis.

## Methods

### Animals and experimental design

This study was approved by the Animal Research Committee of Saitama Prefectural University (approval number: 2020-6). The animals were handled according to relevant legislation and institutional guidelines for humane animal treatment. Here, 18 C57BL/6 male mice were used to evaluate the methods used to create novel non-invasive ACL-R models and to determine whether any intra-articular tissue injuries were induced. The contralateral knee joint was used in the INTACT group (Fig. 1A). A total of 54 mice were randomly classified into three groups: ACL-R group (n = 18; joint instability increased by ACL rupture), CATT group (n = 18; joint instability with ACL rupture suppressed by external support), and Sham group (n = 18). Six mice in each group were sacrificed at 2, 4, and 8 weeks, and the target tissues were collected for each analysis (Fig. 1B). All mice were housed in plastic cages under a 12-hour light/dark cycle. The mice were allowed unrestricted movement within the cage and had free access to food and water.

**Figure 1.**
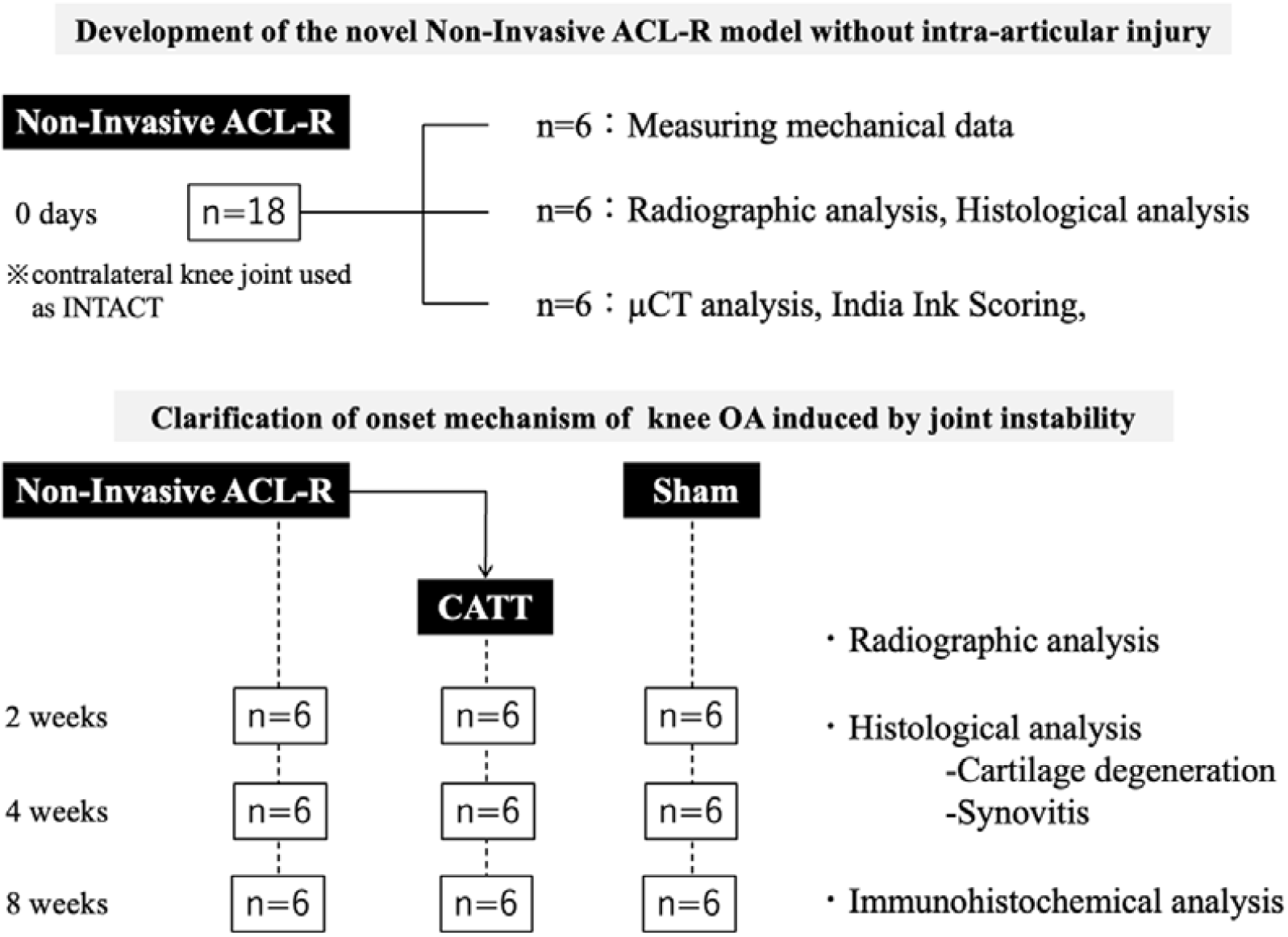
Experimental design. (A) Development of the novel non-Invasive ACL-R model without intra-articular injury. We created the non-Invasive ACL-R model and performed a biomechanical investigation and histological, morphological, and macroscopical analyses to evaluate the intra-articular injuries, except for ACL rupture. (B) Clarification of the onset mechanism of knee OA induced by joint instability. Non-Invasive ACL-R group, CATT group, and Sham group mice were sacrificed at 2, 4, and 8 weeks and the degree of OA progression was assessed through validity investigation and histological analyses.

### Development of a novel non-invasive ACL-R model without intra-articular injury

#### Creating the model

All procedures were performed on the left knee joint of each mouse under a combination of anesthetics drugs (medetomidine, 0.375 mg/kg; midazolam, 2.0 mg/kg; and butorphanol, 2.5 mg/kg). The knee joint was fixed at 90° using surgical tape on a stand. The femoral condyle was pushed manually in the long-axis direction, causing ACL rupture due to the relative anterior dislocation of the tibia. To evaluate the intra-articular injuries other than the ACL injury, we collected the knee joints 0 days after creating the model and performed the following analyses (Fig. 1B).

#### Calculating the force of ACL rupture

To assess the force of ACL rupture, an ACL-R model was created using a load cell device, and mechanical data were collected. The data were transmitted to the PC and processed using a load cell amplifier, Arduino, and Jupyter Lab software to calculate the force and speed of ACL rupture (Fig. 2A). Details were described in the *Supplementary Methods*.

**Figure 2.**
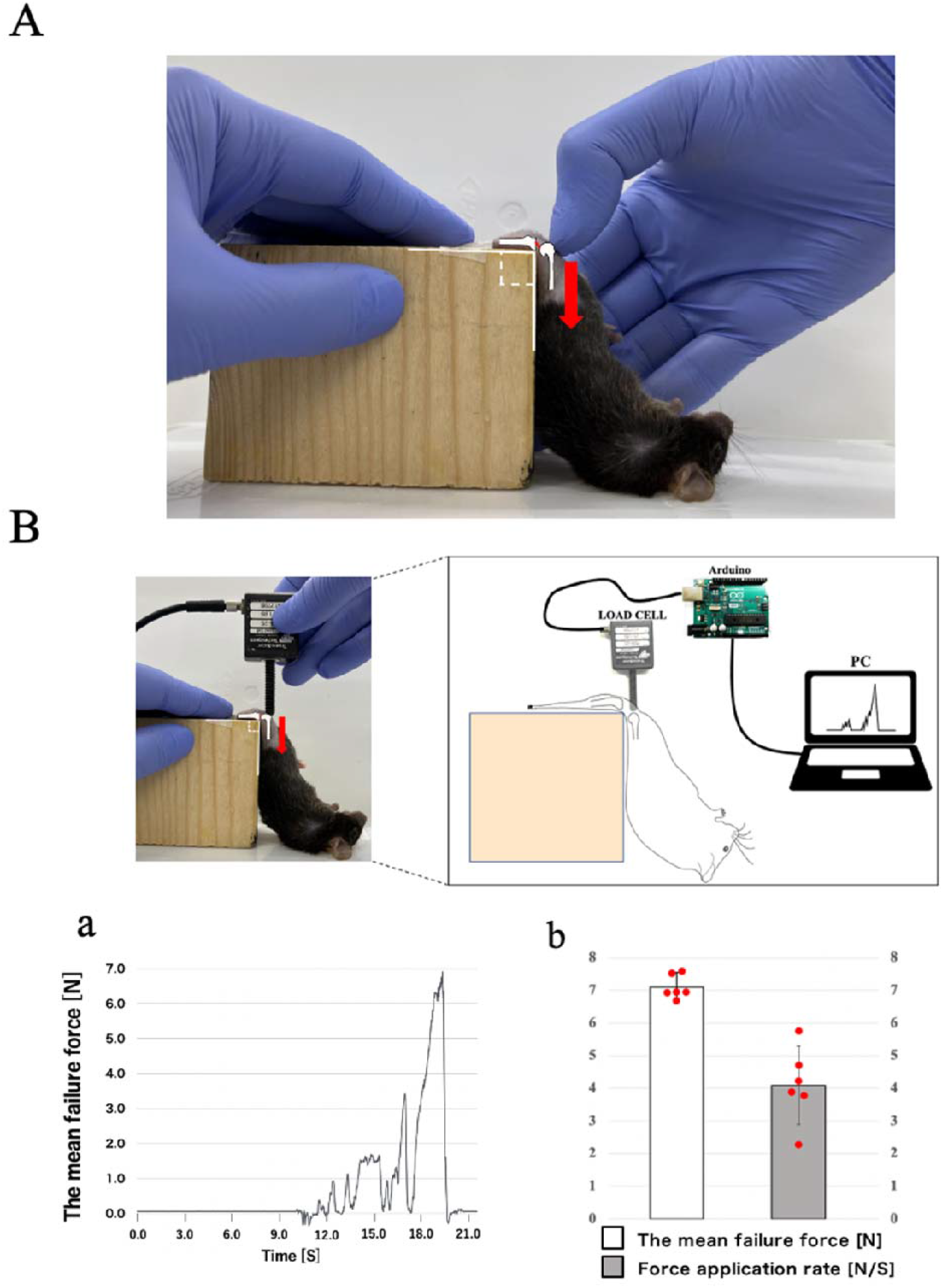
(A) Novel non-invasive ACL-R model. We fixed the knee joints at 90°, pushed the femoral condyle manually in the long axis, and then confirmed ACL rupture and anterior tibial displacement. (B) Mechanical data of ACL rupture. Using MINI LOAD CELL and Arduino, the longitudinal mechanical data was acquired during ACL rupture (a). Then, the mean failure force causing ACL rupture and the force application rate were calculated. Data are presented as the mean ± 95% CI.

#### Radiographic analysis

We performed the anterior drawer test to evaluate the joint instability based on a previous study ^15^. Briefly, the knee joints were fixed at 90 ° flexion position to the device, then the proximal tibia was pulled forward with a constant force spring (0.05 kgf; Sanko Spring Co., Ltd., Fukuoka, Japan). Then, X-ray images were taken, and the anterior displacement was measured by the linear distance from the perpendicular line from the anterior end of the femoral condyle to the tibial tuberosity (ImageJ; National Institutes of Health, Bethesda, MD, USA).

#### Histological analysis

Knee joints were fixed using 4% paraformaldehyde for 1 day, decalcified in 10% ethylenediaminetetraacetic acid for 2 weeks, dehydrated, and embedded in paraffin. The samples were cut in the sagittal plane (thickness, 7 μm) using a microtome. Hematoxylin and eosin (H&E) and safranin-O/fast green stainings were performed to observe ACL rupture and to evaluate the articular cartilage and meniscus injuries, respectively.

#### Micro-computed tomography (μCT) analysis

To evaluate the morphological changes occurring in subchondral bone, the bone volume/tissue volume fraction (BV/TV, %), trabecular thickness (Tb.Th, mm), trabecular number (Tb.N, 1/mm), and trabecular separation (Tb.Sp, mm) in Medial Femoral Condyle (MFC), Lateral Femoral Condyle (LFC), Medial Tibial Plateau (MTP), and Lateral Tibial Plateau (LTP) were determined using CTAn software (BRUKER, MA, USA) ^15^. Details were described in the *Supplementary Methods*. Furthermore, we searched for macroscopic bone loss from the reconstructed image.

#### Macroscopic Observation analysis

After the μCT analysis, the tibial plateau was carefully separated from the femoral condyle, and these tissues were stained with India ink to visualize the damaged region on the joint surface ^19^. A stereomicroscope was used to take macroscopic images of the femoral and tibial joint surfaces. The cartilage lesions of the MFC, LFC, MTP, and LTP were evaluated using a macroscopic score of 0–5 points.

### Clarification of the onset mechanism of knee OA induced by joint instability

#### Creating the model

The non-invasive ACL-R model was created as described above, and the CATT model was created according to a previous study^15)^. Detailed protocols for making the CATT and Sham models are described in the Supplementary Methods. After creating each model, we collected the knee joints at 2, 4, and 8 weeks and performed the following analyses.

#### Radiographic analysis

To evaluate the reproduction of joint instability *in vivo*, anterior drawer test was performed as described above, and the anterior displacement of the tibia was quantified.

#### Histological analysis

We performed safranin-O/fast green staining to histologically evaluate the articular cartilage degeneration and synovitis during OA progression. The Osteoarthritis Research Society International (OARSI) histopathological grading system ^20^ and synovitis score ^21^ were assessed by two independent observers blinded to all other sample information. The contact area not covered by the meniscus on the MTP was used as the OARSI score. The medial synovium located inside the infrapatellar pad was used to determine the synovitis score. The mean of the observer’s scores was used as a representative value.

#### Immunohistochemical analysis

To assess the expression of MMP-3 and tumor necrosis factor-α (TNF-α), immunohistochemical staining was performed using anti-MMP-3 (1:100, ab52915, Abcam) and TNF-α (1:200, bs-1110R, Bioss). Detailed protocols are described in the *Supplementary Methods*. Then, we calculated the ratio between the number of MMP-3- and TNF-α-positive cells and the number of chondrocytes in the articular cartilage area and that of synovial cells in a synovial area of 10,000 μm^2^ (100 μm × 100 μm).

#### Statistical analysis

Statistical analysis of the measured data was performed using R software version 3.6.1. The Shapiro-Wilk test was used to verify the normality of all the analyzed data. We conducted a paired-sample t-test for the anterior drawer test immediately after creating the non-invasive model. For India ink score, BV/TV, Tb.Th, Tb.N, and Tb.Sp, Wilcoxon signed-rank test was performed. One-way analysis of variance was performed for the anterior drawer test at 2, 4, and 8 weeks. For comparing the number of MMP-3- and TNF-α-positive cells in the articular cartilage and synovium, the Tukey-Kramer test was used.

Meanwhile, the Kruskal-Wallis test was used to compare the OARSI and synovitis scores, and the Steel-Dwass method was used for the subsequent multiple comparisons. Parametric data are expressed as the means ± 95% confidence intervals (95% CI), whereas the non-parametric data are expressed as the medians ± interquartile ranges. Statistical significance was set at p < 0.05.

## Results

### Development of the novel Non-Invasive ACL-R model without intra-articular injuries

#### Measurement of the ACL rupture force

To acquire the mechanical data, the ACL rupture force and time were measured using a load cell during the creation of a non-invasive ACL-R model. The mean failure force causing ACL rupture was 7.11 ± 0.45 N, and the time taken for rupture was 1.90 ± 0.79 s; the force application rate derived from these results was 4.09 ± 1.21 N/s (Fig. 2B).

### Anterior tibial displacement was confirmed in the Non-Invasive ACL-R model

Soft X-ray images of the knee joints during the anterior drawer test after creating the non-invasive ACL-R model are shown in Fig. 3A. Compared with the INTACT group, an obvious abnormality in the tibial position was observed, and the anterior tibial displacement was significantly increased in the ACL-R group (p < 0.001, 95% CI = [0.824–1.126]).

**Figure 3.**
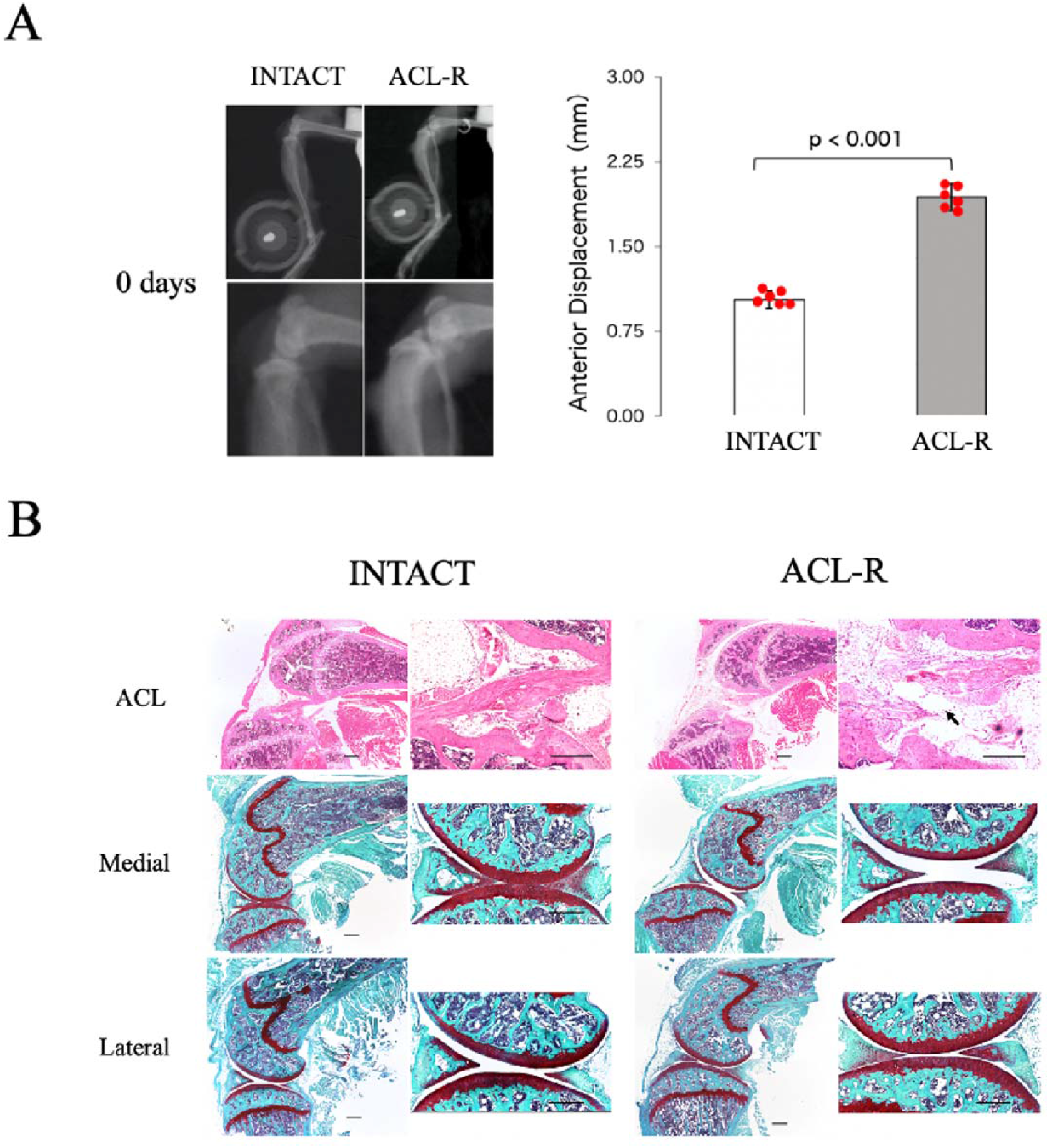
(A) Evaluation of ACL rupture using a soft X-ray device. The anterior tibial displacement was significantly increased in the non-invasive ACL-R group compared to that of the INTACT group. Data are presented as the mean ± 95% CI. (B) Histological analysis of the intra-articular tissues using H&E and safranin-O/fast green staining. In the INTACT group, no injury was detected in ACL, cartilage, or meniscus. In the non-invasive ACL-R group, only the ruptured ACL was confirmed; however, no articular cartilage or meniscus injuries were observed. Black scale bar, 300 μm. Black arrows show an ACL disrupted continuity.

### No soft tissue injuries except for ACL rupture were detected histologically

The histological images of the knee joint after H&E and safranin-O/fast green staining are shown in Fig. 3B. In all mice of the ACL-R group, ACL disrupted continuity, and an abnormal anterior tibial displacement was observed; however, no damage was found in the soft tissues, including the cartilage surface and the meniscus, in the medial and lateral compartment.

### No morphological changes in the whole knee joints and subchondral bone were observed

The 3D reconstruction images of the knee joint obtained via μCT analysis and morphological analysis of the subchondral bone are shown in Fig. 4A. No obvious bone loss or avulsion fracture of the ACL attachment was observed in the ACL-R group. Moreover, there were no significant differences in the BV/TV, Tb.Th, Tb.N, and Tb.Sp in each compartment between the INTACT and ACL-R groups (Detailed results showed in *Supplementary data*).

**Figure 4.**
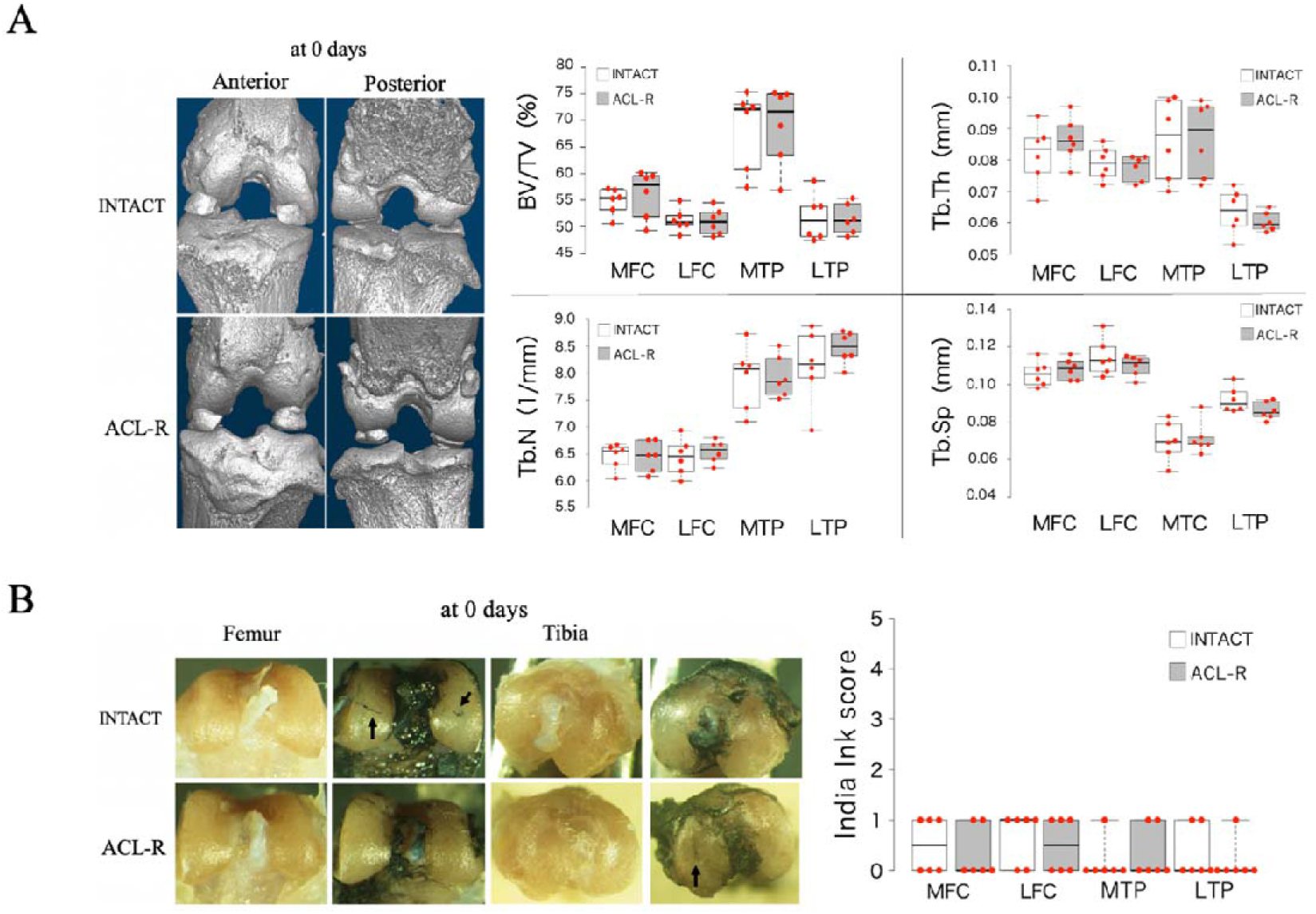
(A) The 3D images of the whole knee joint and morphological analysis of the subchondral bone using μCT. No bone loss or microfracture was observed in the non-invasive ACL-R group. Moreover, there were no differences in the BV/TV, Tb.Th, Tb.N, and Tb.Sp in the MFC, LFC, MTP, and LTP between the INTACT and the ACL-R groups. Data are presented as the median ± interquartile range. (B) The macroscopic images of the joint surface stained with India ink. There were no differences in MFC, LFC, MTP, and LTP between the INTACT and the ACL-R groups. Data are presented as the median ± interquartile range. Black arrows show the punctate depressions.

### No joint surface injuries were detected using India ink staining

The macroscopic images of the articular surface and India ink scoring results are shown in Fig. 4B. Partial staining was observed in each compartment in the INTACT and ACL-R groups; however, no significant differences were observed between the groups for each compartment (Detailed results showed in *Supplementary data*).

### Clarification of the onset mechanism of knee OA induced by joint instability

#### Anterior drawer test

To evaluate the validity of each model, we performed an anterior drawer test (Fig. 5). At 2 weeks, the anterior displacement in the ACL-R group was significantly higher than that in the Sham and CATT groups (ACL-R vs. Sham, p < 0.001; 95% CI = [-1.482 to - 0.897]; ACL-R vs. CATT, p < 0.001; 95% CI = [-1.211 to -0.625]). At 4 weeks, the anterior displacement in the ACL-R group was significantly higher than that in the Sham and CATT groups (ACL-R vs. Sham, p < 0.001; 95% CI = [-1.508 to -0.871]; ACL-R vs. CATT, p < 0.001; 95% CI = [-1.252 to -0.614]). At 8 weeks, the anterior displacement in the ACL-R group was significantly increased compared with that in the Sham and CATT groups (ACL-R vs. Sham; p < 0.001, 95% CI = [-1.509 to -0.966]; ACL-R vs. CATT, p < 0.001, 95% CI = [-0.841 to -0.298]), and the anterior displacement in the CATT group was also significantly increased compared with that in the Sham group (p < 0.001; 95% CI = [-0.939 to -0.396]).

**Figure 5.**
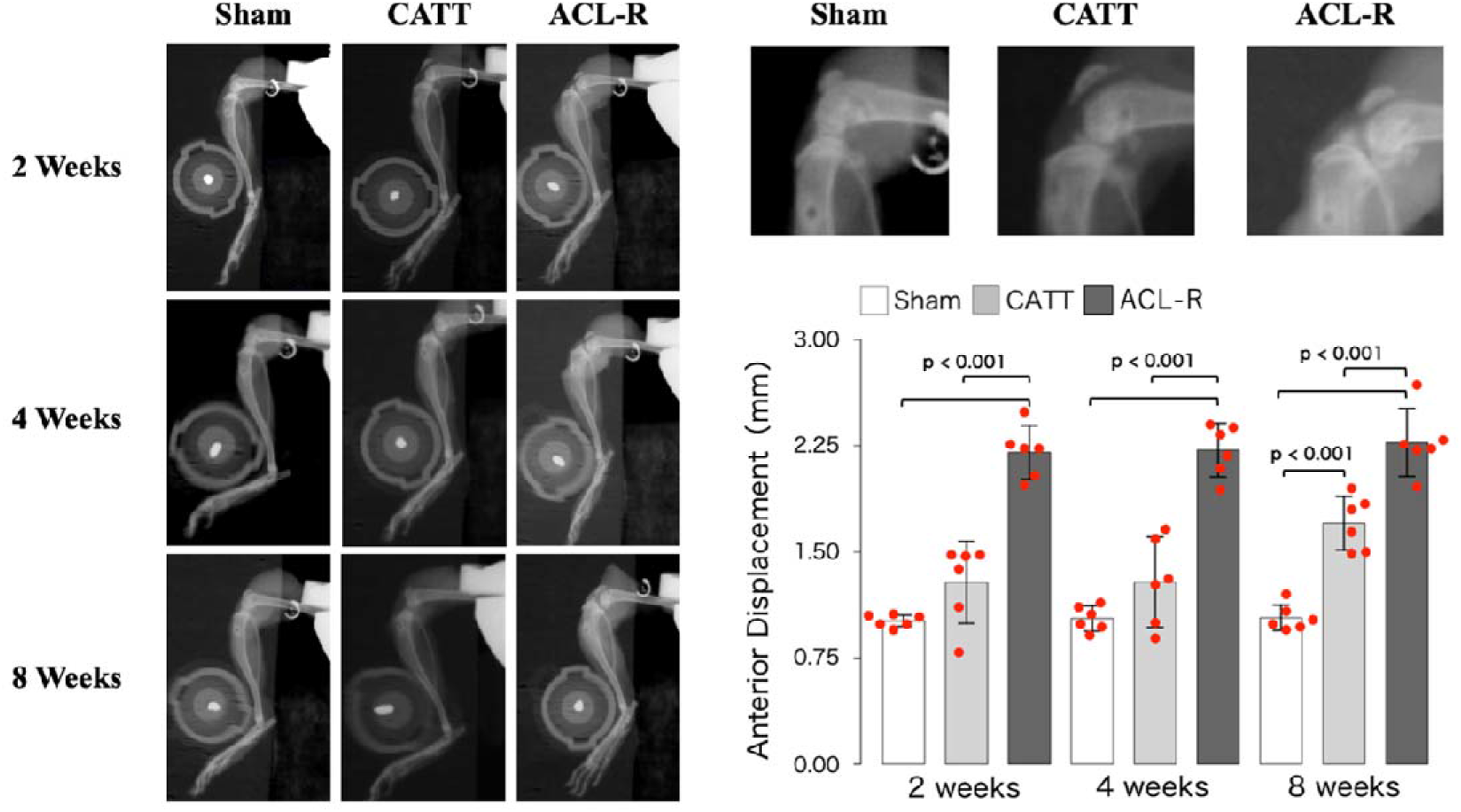
Evaluation of the joint instability *in vivo* using soft X-ray analysis. At 2, 4, and 8 weeks, the amount of anterior tibial displacement was significantly increased in the ACL-R group compared with that in the Sham and CATT groups. Data are presented as the mean ± 95% CI.

### Histological analysis of cartilage degeneration

The safranin-O/fast green staining images and the OARSI score results are shown in Fig. 6A. The safranin-O staining intensity in the surface layer of the articular cartilage decreased in all groups at 2 weeks. Irregularities and fibrillation of the articular cartilage surface were observed in the CATT and ACL-R groups, but not in the Sham group, at 4 weeks. However, there was no significant difference in the OARSI scores at 2 and 4 weeks.

**Figure 6.**
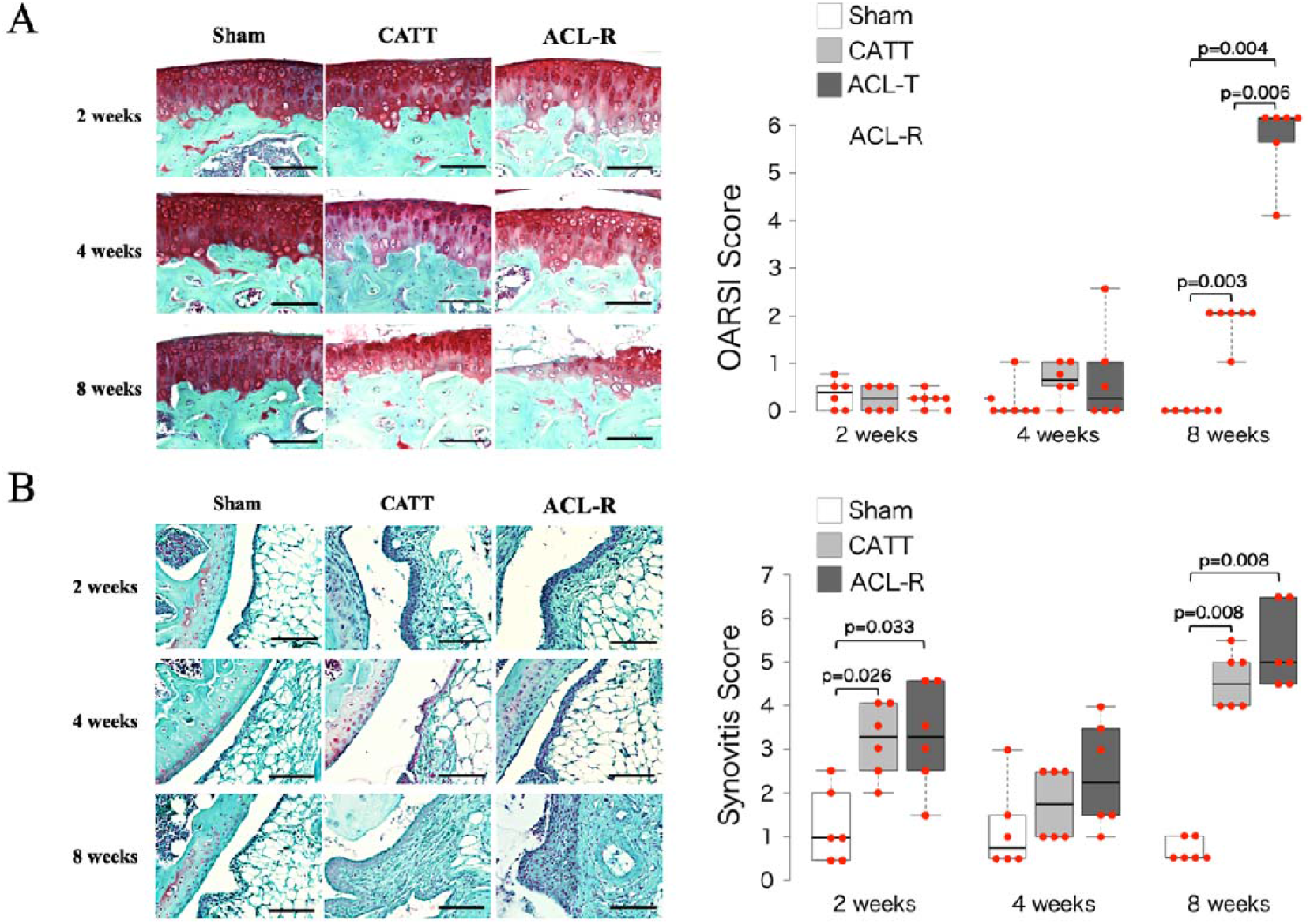
(A) Histological analysis of cartilage degeneration using safranin-O/fast green staining and OARSI score. At 8 weeks, in the CATT group, moderate cartilage degeneration was observed, and the OARSI score was significantly increased compared to that in the Sham group. At 8 weeks, in the ACL-R group, severe cartilage degeneration was observed and the OARSI score was significantly increased compared with that in the Sham and CATT groups. Data are presented as the median ± interquartile range. Black scale bar, 100 μm. (B) Histological analysis of synovitis using safranin-O/fast green staining and synovitis score. At 2 weeks, in the CATT and ACL-R groups, moderate enlargement of the synovial lining cell layer and increased synovial cellularity of the synovial stroma were observed. Furthermore, the synovitis score was significantly increased in these groups compared to that of the Sham group. At 8 weeks, in the CATT and ACL-R groups, severe enlargement of the synovial lining cell layer, increased synovial cellularity of the synovial stroma, and higher inflammatory infiltration were observed. The synovitis score increased significantly compared to that in the Sham group. Data are presented as the median ± interquartile range. Black scale bar, 100 μm.

At 8 weeks, partial clefts and erosion in the surface layer of the articular cartilage were observed in the CATT group; vertical cracks and massive erosion in the calcified cartilage layer were observed and remarkable cartilage degeneration was confirmed in all mice in the ACL-R group. Moreover, the OARSI score in the ACL-R group was significantly higher than that in the Sham and CATT groups (ACL-R vs. Sham, p = 0.004; ACL-R vs. CATT, p = 0.006), and that of the CATT group was significantly increased compared with that of the Sham group (p = 0.003).

### Histological analysis of synovitis

The safranin-O/fast green staining images and synovitis score results are shown in Fig. 6B. At 2 weeks, a moderate enlargement of the synovial lining cell layer and increased cellularity of the synovial stroma were observed in the CATT and ACL-R groups. The synovitis score of the CATT and ACL-R groups was significantly higher than that of the Sham group (CATT vs. Sham, p = 0.026; ACL-R vs. Sham, p = 0.033). However, there was no significant synovitis score difference between the CATT and ACL-R groups.

At 4 weeks, although mild enlargement and increased cellularity were observed in the CATT and ACL-R groups, the synovitis scores weren’t significantly different between these groups. At 8 weeks, the CATT and ACL-R groups showed severe enlargement, increased interstitial cellularity, and inflammatory infiltration. The synovitis score of the CATT and ACL-R groups was significantly higher than that of the Sham group (CATT vs. Sham, p = 0.008; ACL-R vs. Sham, p = 0.008). However, there was no significant synovitis score difference between the CATT and ACL-R groups.

### Immunohistochemical analysis of the articular cartilage and synovium

The immunohistochemical staining of the articular cartilage and the positive cell rate analysis results are shown in Fig. 7. At 2 weeks, there were no significant differences in the number of MMP-3-positive cells between the groups. At 4 weeks, the number of MMP-3-positive cells in the ACL-R group was significantly higher than that in the Sham and CATT groups (ACL-R vs. Sham, p = 0.001; 95% CI = [-37.44, -9.485]; ACL-R vs. CATT, p = 0.043; 95% CI = [-28.32 to -0.364]). At 8 weeks, the number of MMP-3-positive cells in the ACL-R group was significantly higher than that in the Sham and CATT groups (ACL-R vs. Sham, p < 0.001; 95% CI = [-36.363 to -13.986]; ACL-R vs. CATT, p = 0.031; 95% CI = [-23.398 to -1.021]), and that in the CATT group was significantly higher than that in the Sham group (CATT vs. Sham, p = 0.022; 95% CI = [-24.153 to -1.776]) (Fig. 7A). There were no significant differences in the number of TNF-α-positive cells between the groups at 2, 4, and 8 weeks (Fig. 7B).

**Figure 7.**
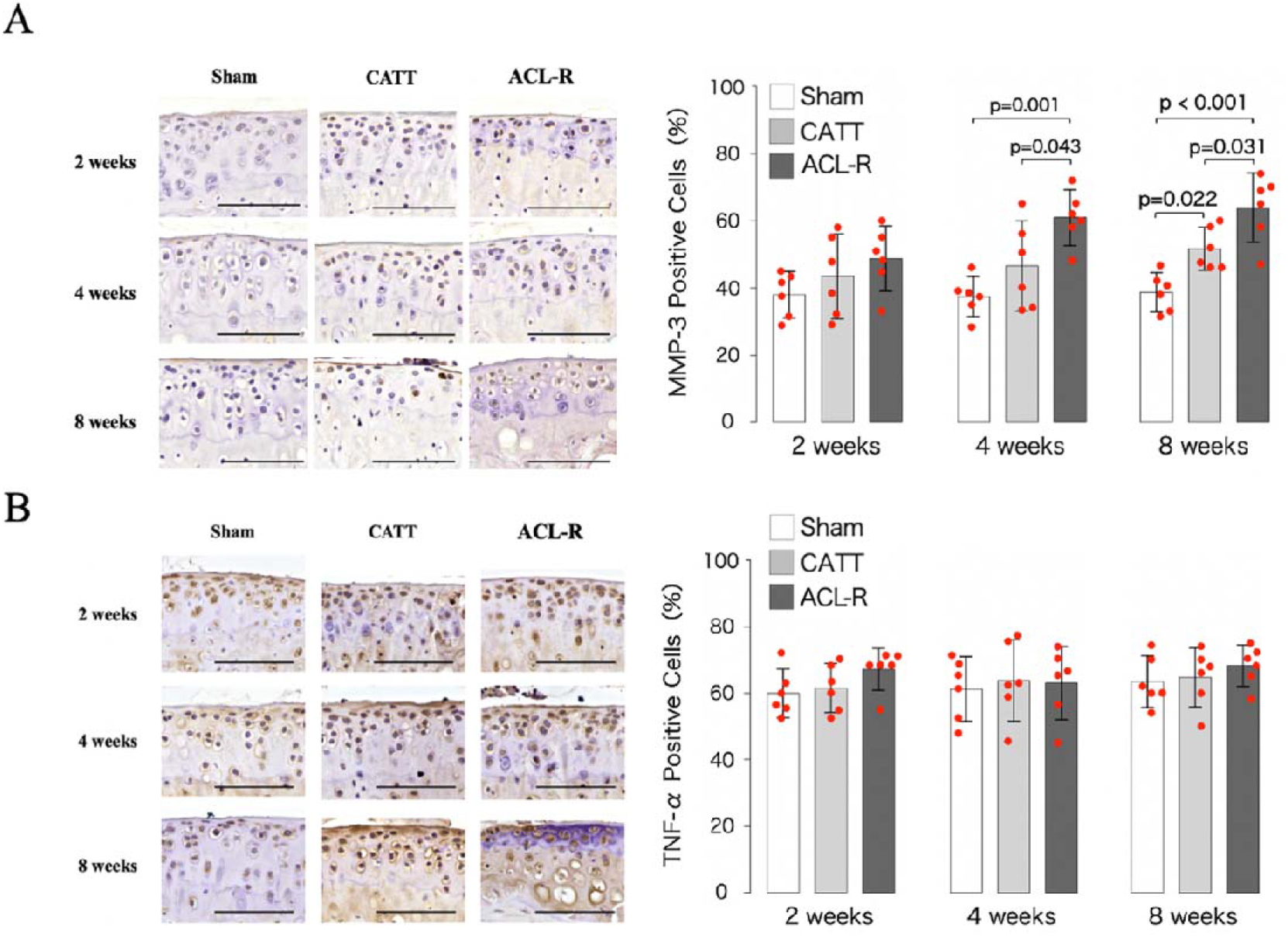
(A) Immunohistochemical analysis of MMP-3 in cartilage. At 4 weeks, the MMP-3-positive cell rate in the ACL-R group was significantly increased compared with that in the Sham and CATT groups. At 8 weeks, the MMP-3-positive cell rate in the ACL-R group was significantly increased compared with that in the Sham and CATT groups, and that in the CATT group was significantly increased compared with that in the Sham group. Data are presented as the mean ± 95% CI. Black scale bar, 50 μm. (B) Immunohistochemical analysis of TNF-α in cartilage. At 2, 4, and 8 weeks, there were no differences in the TNF-α-positive cell rate among all groups. Data are presented as the mean ± 95% CI. Black scale bar, 50 μm.

The immunohistochemical staining in the synovium and the positive cell rate analysis results are shown in Fig. 8. At 2 weeks, there were no significant differences in the number of MMP-3-positive cells between the groups. At 4 weeks, the number of MMP-3-positive cells in the ACL-R group was significantly higher than that in the Sham group (p = 0.014; 95% CI = [-27.612 to -3.01]). At 8 weeks, the number of MMP-3-positive cells in the ACL-R group was significantly higher than that in the Sham and CATT groups (ACL-R vs. Sham, p < 0.001; 95% CI = [-34.197 to -15.782]; ACL-R vs. CATT, p = 0.042; 95% CI = [-18.697 to -0.282]), and that in the CATT group was significantly higher than that in the Sham group (CATT vs. Sham, p = 0.001; 95% CI = [-24.707 to -6.292]) (Fig. 8A). There was no significant difference in the number of TNF-α-positive cells at 2 weeks between the groups; however, that of the ACL-R group was the highest and that of the CATT group was higher than that of the Sham group. At 4 weeks, there was no significant difference in the number of TNF-α-positive cells between the groups. At 8 weeks, the number of TNF-α-positive cells in the CATT and ACL-R groups was significantly higher than that in the Sham group (CATT vs. Sham, p = 0.032; 95% CI = [-31.833 to -1.354]; ACL-R vs. Sham, p = 0.011; 95% CI = [-34.826 to -4.347]) (Fig. 8B).

**Figure 8.**
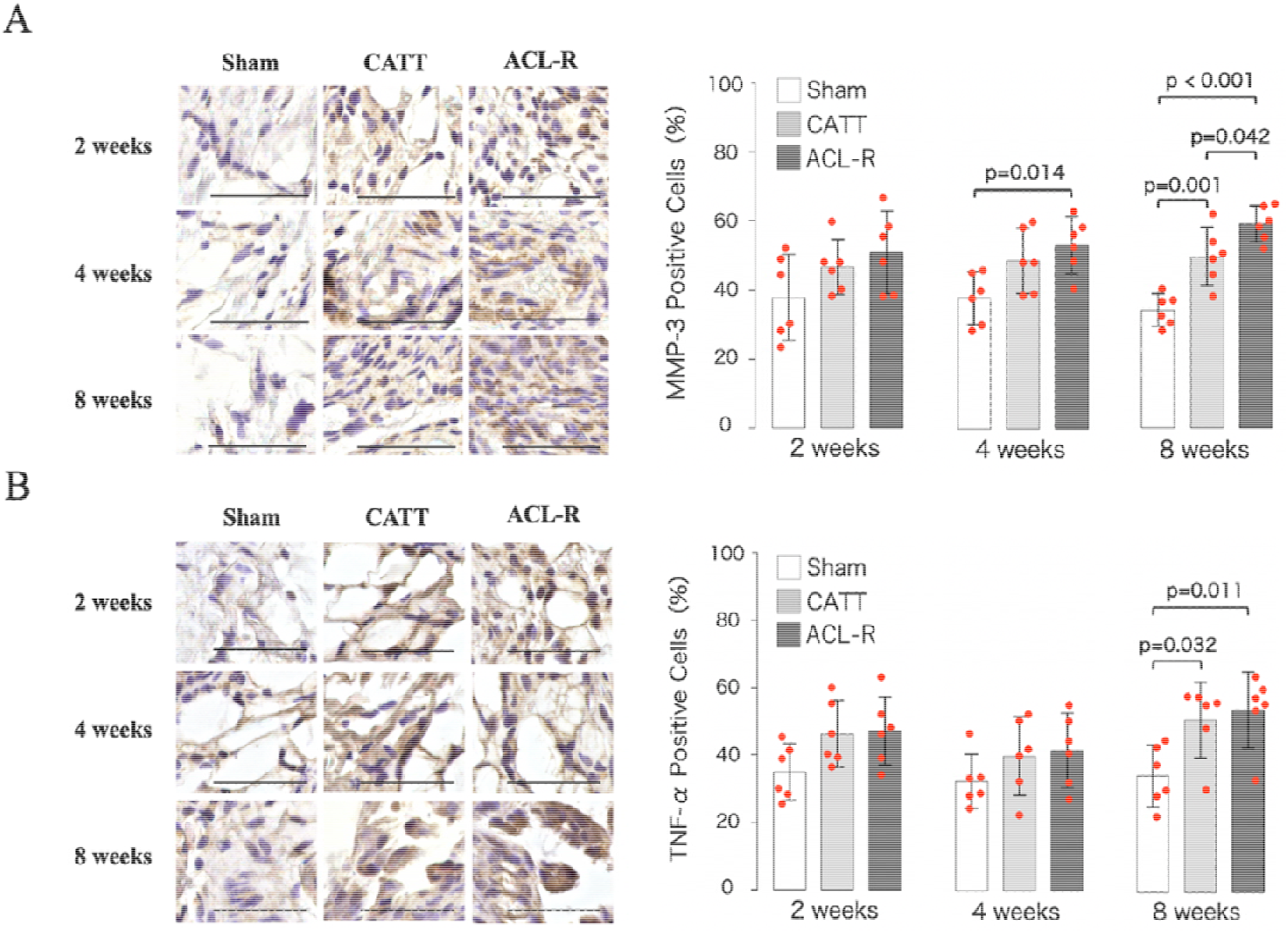
(A) Immunohistochemical analysis of MMP-3 in the synovium. At 4 weeks, the MMP-3-positive cell rate in the ACL-R group was significantly increased compared with that in the Sham group. At 8 weeks, the MMP-3-positive cell rate in the ACL-R group was significantly increased compared with that in the Sham and CATT groups, and that in the CATT group was significantly increased compared with that in the Sham group. Data are presented as the mean ± 95% CI. Black scale bar, 50 μm. (B) Immunohistochemical analysis of TNF-α in the synovium. At 8 weeks, the TNF-α-positive cell rate in the CATT and ACL-R groups was significantly increased compared to that in the Sham group. Data are presented as the mean ± 95% CI. Black scale bar, 50 μm.

## Discussion

We aimed to establish a novel non-invasive ACL-R mouse model without intra-articular tissue injuries and to elucidate the effect of anterior-posterior joint instability on the onset mechanism of knee OA. Our ACL-R model showed no injuries including cartilage degeneration, meniscus lesions, and subchondral bone loss. Then, we evaluated knee OA progression using ACL-R and CATT models, which suppresses joint instability. The CATT group showed reduced cartilage degeneration and MMP-3 expression in the articular cartilage and synovium compared to those of the ACL-R group. However, there were no differences in synovitis and the TNF-α levels in the articular cartilage and synovium between the CATT and ACL-R groups.

In the novel non-invasive ACL-R group, ACL rupture was observed histologically, and the anterior tibial displacement was significantly increased compared with that in the INTACT group. A previous study using C57BL/6 mice revealed that the mean ACL rupture force was 6.78 ± 1.53 N and the failure speed was 10.90 ± 4.37 N/s ^22^. Our data support these rupture force results, which may be due to the fact that a similar method was used to push the femoral condyle in the long-axis direction. Conversely, other reports showed that a force of 12 N was required for ACL rupture in C57BL/6 mice using a compression device ^17, 18^. We obtained a lower force value than the previous study using a compression device, which may be attributed to differences in the knee flexion degree and contact of the joint surfaces ^23^. No histological, macroscopic, or morphological differences were found between the INTACT and ACL-R groups.

Cartilage and meniscus injuries, and subchondral fractures occur in half, more than half, and 80%–90% of ACL injuries, respectively ^24^. These injuries are attributed to compression stress induced by the contact between the femoral and tibial surfaces, resulting in the apoptosis of chondrocytes, weakening of the load distribution due to meniscal dysfunction, and changes in bone remodeling, leading to knee OA. Considering the alignment of the femur and tibia in the conventional non-invasive ACL rupture model^16-18^, it can be estimated that the rupture force is applied in the direction of compression of the joint surfaces. In such cases, the effects of compression force and joint instability may be mixed in OA progression. For these reasons, we established a novel non-invasive ACL-R model which less likely to cause compression stress to evaluate joint instability. This model enables us to assess the effect of joint instability on OA progression, which is frequently observed in knee OA patient with laxity^25-27^. Moreover, ACL suppresses the shear-stress in the knee joint^28^; therefore, it is perhaps possible to evaluate the effects of shear-stress on OA progression.

Then, we examined the effects of joint instability on OA progression. Our previous study has already revealed that anterior-instability by ACL dysfunction probably causes degeneration of surfaces cartilage before subchondral bone changes measured by μCT ^15^. Therefore, this study focused on analyzing cartilage degeneration on the joint surface.The histological results showed that the OARSI score in the ACL-R group was significantly higher than that in the CATT group at 8 weeks. The factors that cause knee OA after ACL injury include the acute effects of intra-articular damage and the chronic mechanical effect caused by kinematic changes ^29^. The ACL-R model used here is confirmed to cause no injuries. Whereas the anterior tibial displacement in the ACL-R group was significantly increased compared with that of the Sham and CATT groups. Therefore, cartilage degeneration in the ACL-R group at 8 weeks was most likely caused by abnormal mechanical stress. MMP-3 is a protease that triggers the expression of other MMP family members by being expressed early in knee OA and is correlated with the severity of knee OA ^3, 30^. Additionally, chondrocytes and synovial cells respond to shear-stress and promote MMP-3 expression ^31, 32^. Here, the number of MMP-3-positive cells among the chondrocytes significantly increased 1.3-fold at 4 weeks in the ACL-R group compared with that in the CATT group. The number of MMP-3-positive cells among the chondrocytes and synovial cells significantly increased 1.23-fold and 1.19-fold at 8 weeks in the ACL-R group compared to that in the CATT group, respectively. These results suggest that chondrocytes are the first cells to respond to anterior-posterior joint instability *in vivo*, causing cartilage degeneration.

Synovitis predates cartilage degeneration and is thought to be the source of intra-articular degeneration by propagating catabolic factors to chondrocytes via the synovial fluid ^10, 12^. Lifan *et al*. reported that synovitis occurred as early as 1 week after DMM, which precedes cartilage degradation, subchondral sclerosis, and osteophyte formation ^11^. However, the relationship between cartilage degeneration and synovitis under mechanical stress hasn’t been revealed clearly since synovial invasive animal models have been used in these studies. Interestingly, the synovitis scores in the CATT and ACL-R groups were significantly higher than those in the Sham group at weeks 2 and 8, however, there was no difference between the CATT and ACL-R groups. Moreover, although the number of TNF-α-positive cells in the synovium in the CATT and ACL-R groups increased at 2 weeks and significantly increased at 8 weeks compared with those of the Sham group, no difference was observed between the CATT and ACL-R groups. TNF-α alters synovial cells, inducing an inflammatory phenotype, and is released into the synovial fluid soon after ACL injury ^33^. Therefore, synovitis in the CATT and ACL-R groups at 2 weeks may indicate that the acute inflammation associated with ACL rupture affected the synovial membrane through the synovial fluid. Chronic synovitis in knee OA generally results from an innate immune mechanism mediated by Damage-associated molecular patterns (DAMPs) in synovial fluid^34^. Considering that there was no difference in the synovitis score between the CATT and ACL-R groups at 8 weeks in which both groups developed cartilage degeneration, the synovitis may be the secondary result responding to the release of DAMPs into the synovial fluid with OA progression.

This study has three limitations: first, while creating our model is relatively easy, it could be subject to individual user competency for causing no intra-articular injuries.

The second is the inability to quantify joint instability occurring *in vivo* in shear-stress. To compensate for the result of this study, *in vitro* experiments that can reliably provide shear-stress are needed in the future. The last one is that no data to support our hypothesis about the onset mechanism of synovitis was provided; therefore, additional experiments about innate immunity, such as DAMPs, need to be conducted.

In conclusion, we successfully established a new non-invasive ACL-R model without intra-articular injuries, in which knee OA is induced by anterior-posterior joint instability. In joint instability-induced OA progression, chondrocytes first show a molecular biological response, leading to a local increase in the MMP-3 level.

Subsequently, the MMP-3 expression may be increased in synovial cells through molecular biological interactions. Our results also suggest that mechanical stress doesn’t directly induce synovitis, which may be indirectly caused by the intra-articular degeneration associated with knee OA progression.

## Acknowledgments

The author(s) received no financial support for the research, authorship, and/or publication of this article. And we would like to thank Editage (www.editage.com) for English language editing.

## Author contributions

All authors approved the final submitted manuscript.

Study design: KT, TK.

Data collection, Mechanical analysis: KN, RS. Histological analysis: KT, KA, SE, YU, KO, HT, and HT. Morphological analysis: KA Manuscript composition: KA, TK.

