## Supplementary data for "Joint Instability Causes Catabolic Enzyme Production in Chondrocytes prior to Synovial Cells in Novel Non-Invasive ACL ruptured Mouse Model"

**Methods**

**Calculating the force of ACL rupture**

The MDB-25 load cell (Transducer Techniques, CA, USA) and load cell amplifier - HX711 (Sparkfun, CO, USA) were used to calculate the ACL rupture force. Concretely, offset adjustment for the load cell was performed using a 30.7271 g weight, the correction function was obtained, and the voltage values were calibrated. After these procedures, a load cell amplifier was connected to the PC, and we opened the Arduino and Jupyter Lab software on the PC. Then, the mouse femoral condyle was pushed in the long axis using a bolt attached to a load cell. The mean failure force and the force application rate were calculated based on the mechanical data measured by a load cell.

**Micro-computed tomography (µCT) analysis**

For analysis of the subchondral bone, the knee joints were scanned using a microCT system (Skyscan 1272, BRUKER, MA, USA) with the following parameters: pixel size, 6 μm; voltage, 60 kV; current, 165 mA. Subsequently, the reconstructed image was acquired using the NRecon software (BRUKER, MA, USA). We designated the region of interest on 40 slides with a 0.7mm circle of the subchondral bone in each region. Considering the stress on the joint surface that may occur when an ACL rupture was induced at 90 degrees of knee flexion, the analysis area was middle in the MFC and LFC, whereas that was posterior in MTP and LTP (Supplementary figure 1). We then calculated BV/TV (%), Tb.Th (mm), Tb.N (1/mm), and Tb.Sp (mm) using CTAn software (BRUKER, MA, USA).

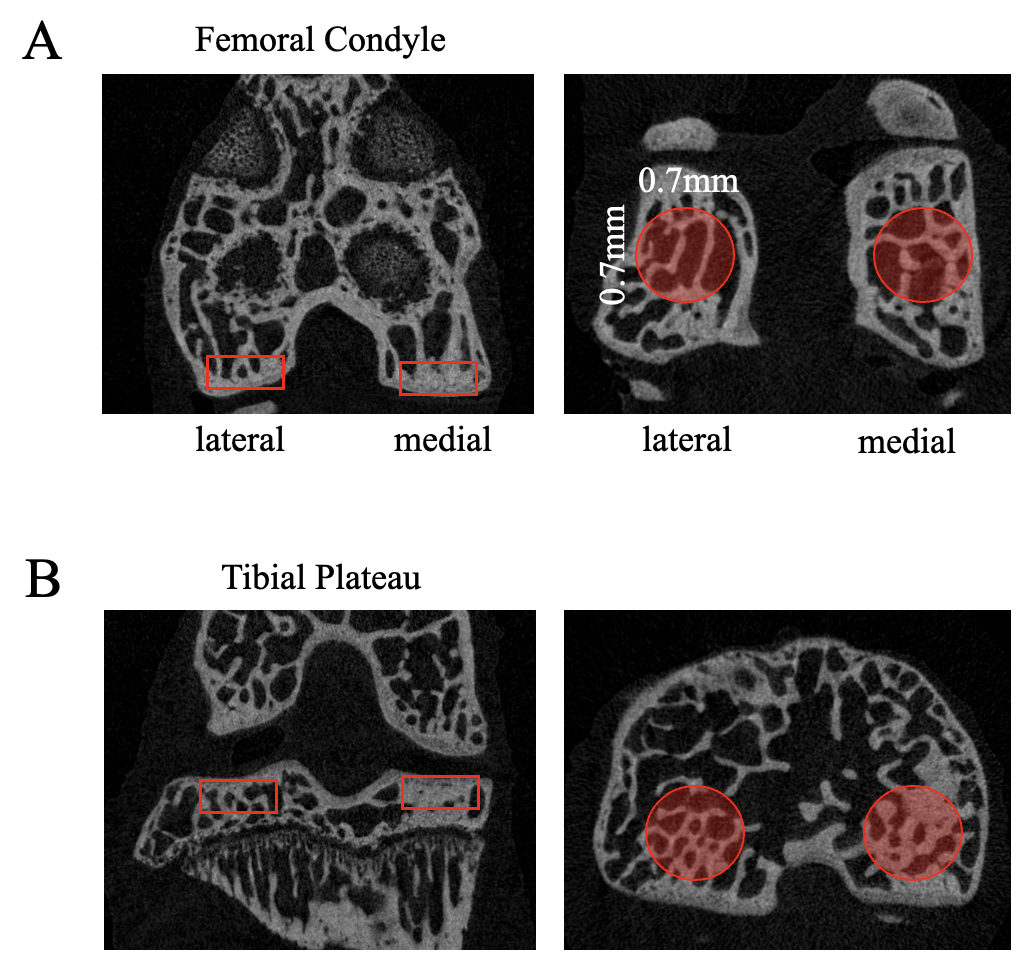

Supplementary figure 1.

The volume of interest in subchondral bone for each region.

A: The subchondral bone inside the medial and lateral femoral condyle.

B: The subchondral bone inside the medial and lateral tibial plateau. All region of interest for each slide was a circle with a diameter of 0.7 mm

**Creating the CATT and Sham model.**

Next, we made the CATT model. After creating the ACL-R model, we made extra-articular bone tunnels in the distal femur and proximal tibia using a 26-gauge needle in the ACL-R model. Then, 4-0 nylon sutures were threaded through these, and the forward displacement of the tibia was suppressed from outside the joint capsule by tying the nylon sutures (Supplementary figure 2).

To standardize the model preparation conditions, bone tunnels were created in the ACL-R groups, and the nylon threads were loosely tied to prevent joint suppression. Finally, we made the Sham model: only extra-articular bone tunnels were created, and the nylon threads were loosely tied.

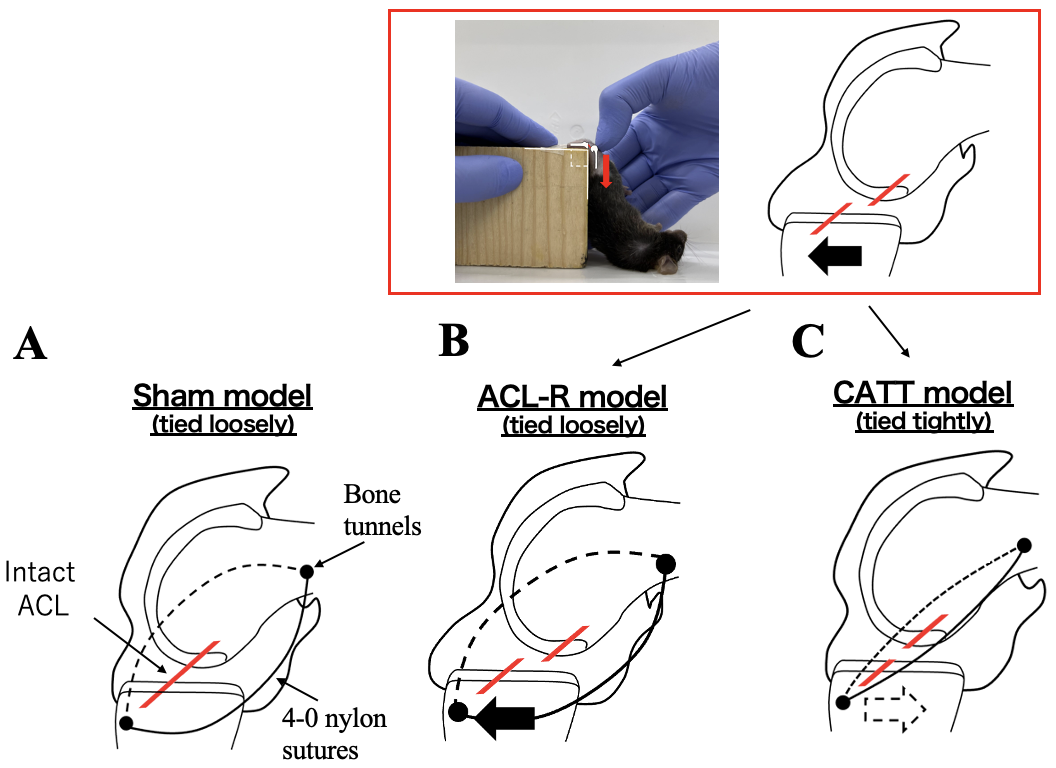

Supplementary figure 2.

The procedures of creating the Sham, ACL-R, and CATT models. A: Sham model. Only extra-articular bone tunnels were made in the distal femur and proximal tibia, and the nylon threads were loosely tied. B: ACL-R model. After inducing ACL rupture, extra-articular bone tunnels were created, and the nylon threads were loosely tied. C: CATT model. After completing the ACL-R model, we made extra-articular bone tunnels, and the nylon threads were tied to suppress anterior tibial translation.

**Immunohistochemical analysis**

To assess the expression of matrix metalloproteinase-3 (MMP-3) and tumor necrosis factor-α (TNF-α), immunohistochemical staining was performed using the avidin-biotinylated enzyme complex method and a Vectastain Elite ABC Rabbit IgG Kit (Vector Laboratories, Burlingame, CA, USA). We used anti-MMP-3 (1:100, ab52915, Abcam) and TNF-α (1:200, bs-1110R, Bioss) as the primary antibodies and anti-rabbit IgG antibody as the secondary antibody. The sections were stained using a Dako Liquid DAB Substrate-Chromogen System (Dako, Glostrup, Denmark). The cell nuclei were stained using hematoxylin at a concentration of 25%.

We calculated the ratio between the number of MMP-3- and TNF-α-positive cells and the number of chondrocytes in the articular cartilage area and that of synovial cells in a synovial area of 10,000 µm2 (100 µm × 100 µm).

**Results**

**No morphological changes in the whole knee joints and subchondral bone were observed**

There were no significant differences in the BV/TV, Tb.Th, Tb.N, and Tb.Sp in each compartment between the INTACT and ACL-R group (BV/TV; MFC; p = 0.437, 95%CI = [-3.990 to 6.015], LFC; p = 0.843, 95%CI = [-1.40 to 2.52], MTP; p = 0.843, 95%CI = [-2.71 to 3.56], LTP; p = 0.843, 95%CI = [-7.18 to 4.87], Tb.Th; MFC; p = 0.156, 95%CI = [-0.011 to 0.007], LFC; p = 0.25, 95%CI = [-0.001 to 0.004], MTP; p = 1, 95%CI = [-0.006 to 0.003], LTP; p = 0.218, 95%CI = [-0.006 to 0.010], Tb.N; MFC; p = 1, 95%CI = [-0.714 to 0.540], LFC; p = 0.562, 95%CI = [-0.500 to 0.404], MTP; p = 843, 95%CI = [-0.503 to 0.495], LTP; p = 0.687, 95%CI = [-1.791 to 0.367], Tb.Sp; MFC; p = 0.312, 95%CI = [-0.009 to 0.006], LFC; p = 0.437, 95%CI = [-0.010 to 0.016], MTP; p = 0.562, 95%CI = [-0.009 to 0.011],LTP; p = 0.218, 95%CI = [-0.006 to 0.023])

**No joint surface injuries were detected using India ink staining**

There were no significant differences in each compartment between the INTACT and ACL-R group (MFC; p = 1, 95%CI = [1 to 1], LFC; p = 1, 95%CI = [1 to 1], MTP; p = 1, 95%CI = [-1 to -1], LTP; p = 1, 95%CI = [1 to 1]).
